# Developmental changes in connectivity between the amygdala subnuclei and occipitotemporal cortex

**DOI:** 10.1101/636894

**Authors:** Heather A. Hansen, Zeynep M. Saygin

## Abstract

The amygdala, a subcortical structure known for social and emotional processing, can be subdivided into multiple nuclei with unique functions and connectivity patterns. Tracer studies in adult macaques have shown that the lateral and basal amygdala subnuclei decrease in connectivity to visual cortical areas moving from anterior to posterior, and that infants have similar adult-like projections plus additional connections that are refined with development. Can we delineate the connectivity between the amygdala subnuclei and occipitotemporal cortex in humans, and will it show similar developmental differences as macaques? If so, what functional regions may be contributing to this pattern of connectivity? To address these questions, we anatomically defined the lateral and basal amygdala subnuclei in 20 adult subjects, 27 kids (aged 7-8), and 15 neonates. We then defined the occipitotemporal region in each individual’s native anatomy, and split this entire region into five equal sections from anterior to posterior. We also defined visual functional parcellations in the occipitotemporal cortex (e.g. FFA, PPA) and anatomically defined primary visual cortex (i.e., V1). Using Diffusion Weighted Imaging data, we ran probabilistic tractography with FSL between the amygdala subnuclei as seeds and the occipitotemporal cortical parcellations as targets. Results showed that like macaques, the mean connectivity across subjects to the occipitotemporal cortex significantly decreased on a gradient from anterior to posterior, and that connectivity in kids and neonates was adult-like but became more refined across development. Further, refinement of connectivity to mid and posterior occipitotemporal cortex was largely driven by anterior PPA, LO, and V1, with connectivity to higher order visual areas increasing with age. The functional maturation of these regions may contribute to the continued refinement of these connections, in line with Interactive Specialization hypotheses of brain development.

## 1. Introduction

How does emotional valence influence visual perception? Whether it be driving by an emotionally salient car crash or happening upon an animal carcass in the jungle, perceiving visual stimuli through an emotional lens can be critical for quick motor responses and ultimate survival. Emotionally salient cues preceding a target actually can enhance target perception (e.g., Phelps et al., 2006), and perceiving aversive stimuli enhances blood flow to the middle temporal gyrus, for example (e.g., Kosslyn et al., 1996). Developmentally, not only does visual acuity improve with age (Teller et al., 1986), but visual perceptual mechanisms of emotional stimuli are also fine-tuned with experience (e.g., Leppänen & Nelson, 2009).

Emotional valence is canonically tied to the amygdala, an evolutionarily preserved neural structure known for emotional processing and regulation (e.g., Ochsner, Silvers, & Buhle, 2012; Phillips et al., 2003). The amygdala has been additionally implicated in social cognition and attention (e.g., Adolphs & Spezio, 2006), fear recognition and conditioning (e.g., Adolphs et al., 2005; Phillips & LeDoux, 1992), stimulus-value learning and reward (e.g., Baxter & Murray, 2002; Paton et al., 2006), and novelty detection (e.g., Kiehl et al., 2005; Schwartz et al., 2003). The functions of the amygdala and the way in which the amygdala assigns valence to stimuli change across development (Tottenham, Hare, & Casey, 2009). Similarly, visual perceptual skills and their neural correlates also change across development (Atkinson, 2000). Perceiving the identity of visual stimuli is commonly attributed to the occipitotemporal cortex, the location of the ventral visual stream and “what” pathway (e.g., Goodale & Milner, 1992). It is posited that emotionally enhanced visual perception may occur via cortical feedback connections between the amygdala and visual cortex (Vuilleumier, 2005).

Work in macaques shows that visual cortical areas that are more rostral receive heavier amygdalar projections than visual cortical areas that are more caudal (Amaral & Price, 1984; Iwai & Yukie, 1987). Projections from the amygdala subnuclei to the ventral visual stream are topographically organized on a gradient, such that more rostral subnuclei project more strongly to rostral visual areas and more caudal subnuclei project more strongly to caudal visual areas (Amaral & Price, 1984; Amaral, 2002; Amaral, Behniea, & Kelly, 2003; Freese & Amaral, 2005, Figure 1). Amaral and colleagues found the basal subnucleus of the amygdala to especially follow this pattern, but noted additional projections from area TE to the lateral subnucleus that creates a feedforward/feedback loop. Other work in adult macaques has similarly shown projections from areas TEO and TE to the lateral nucleus of the amygdala, and from area TE to the basal nucleus (Webster et al., 1991a).

**Figure 1.**
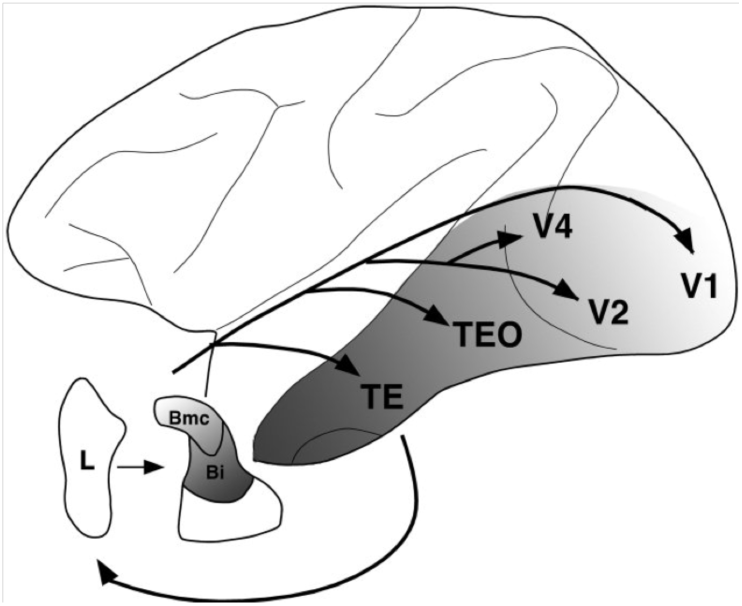
Line drawing summarizing the organization of projections between the amygdala and ventral stream visual cortical areas in adult macaques (reproduced from Freese & Amaral, 2005, pending approval)

Interestingly, these connections change over development. Experiments comparing adult to juvenile animals, specifically in nonhuman primates (e.g., Webster et al., 1991a; Webster et al., 1991b; Kalin et al., 2001; Kalin et al., 1991) and rats (e.g., Bouwmeester et al., 2002a; Bouwmeester et al., 2002b), reveal that amygdalar projections are adult-like in juveniles, but that juveniles also have additional connections that are either totally eliminated with maturation or more refined in their distribution.

Do these connections show a similar pattern in human development? We know that macaque cortex is oriented differently than human cortex, and although homologies exist, the connectivity pattern in macaques may not necessarily perfectly map to humans (Passingham, 2009; Van Essen et al., 2016). Moreover, in humans, it is more challenging to study amygdalar connections, especially with respect to the basal vs. lateral nucleus and their connections to visual cortex. As tracer studies are invasive, studies exploring the structural connectivity of the amygdala and its developmental changes in humans have only been possible more recently, using non-invasive methods such as Diffusion Weighted Imaging (DWI) data. Several groups have used a variety of methods to parcellate the amygdala into two to four subunits (e.g., Saygin et al. 2011; Bach et al., 2011; Brown et al., 2014; Qin et al., 2012; Qin et al., 2014; Gabard-Durnam et al., 2014; Gabard-Durnam et al., 2018). More recent work has made it possible to use local intensity differences in a typical T1 scan to divide the human amygdala into nine separate subunits (Saygin et al., 2017), thus allowing a way to parcellate the amygdala using a T1 and explore the connectivity of these subunits with a separate (independent) connectivity scan. It is advantageous to parcellate the amygdala into its subnuclei because they are functionally distinct. For instance, the lateral subnucleus has been shown to be involved in learning new stimulus-affect associations in humans (Johansen et al., 2010), whereas the basal subnucleus may be the site of convergence of sensory and cognitive information (Stefanacci & Amaral, 2002).

There is some previous work in humans that explores the developmental changes of amygdalar connectivity. A study that explored a cross-sectional sample of 5-30 year olds showed that DWI connectivity of the lateral and basal nuclei to cortical areas becomes increasingly sparse and localized with age (Saygin et al., 2015). A functional connectivity study in 7-9 year olds vs. adults found that the basolateral amygdala had stronger connectivity with temporal regions than the centromedial amygdala, and that overall connectivity was stronger in adults compared to children (Qin et al., 2012). Another study showed that basolateral functional connectivity to regions including parahippocampal gyrus, superior temporal cortex, and occipital lobe decreases with age across 4-23-year-olds (Gabard-Durnam et al., 2014), but that the basolateral amygdala showed increasing functional connectivity to occipital cortex between ages 3 months to 5 years Gabard-Durnam et al., 2018). This last study is the opposite pattern than what was found in macaque development (i.e., decreasing connectivity across age, e.g., Webster et al., 1991a), and may be due to the differences between functional vs. white-matter connectivity or due to differences between macaques and humans. Moreover, it remains unclear why these connections change with development; it is possible that they change with respect to functionally specific parts of occipitotemporal cortex that show increasing developmental specialization. To date, no study has investigated neonatal structural connectivity of the amygdala subnuclei and no study has investigated this connectivity with respect to functional regions within visual cortex.

Does the rostrocaudal gradient of connectivity from the lateral and basal subnuclei observed by Amaral and colleagues in macaques match that of humans, or will a different pattern emerge? More specifically, can we delineate the connectivity between the amygdala subnuclei and the occipitotemporal cortex using noninvasive methods in humans, and does the connectivity differ across development? Moreover, the occipitotemporal cortex contains a multitude of well-studied functional areas in the ventral visual stream; are the developmental changes in connectivity specific to certain functional parcels located within the occipitotemporal cortex? Here we investigate the developmental changes in the connectivity between the lateral and basal amygdalar subnuclei and the occipitotemporal cortex using DWI probabilistic tractography on a cross-sectional sample of adults, kids, and neonates. In Experiment 1, we target the entire cortex anatomically to noninvasively recreate the tracer work done in macaques. In Experiment 2, we apply a unique approach by targeting functionally defined parcels in the ventral visual stream in order to draw conclusions about what might be driving the observed pattern of structural connectivity. To get an even finer-grained idea of the connectivity patterns, we target both the whole parcels and the parcels subdivided into anterior and posterior sections.

## 2. Experiment 1

### 2.1 Method

#### 2.1.1 Participants

##### Adults

Twenty adults (10 female, mean age = 24.6 years) from the greater Boston area were used as part of a larger study exploring the development of the visual word form area. Participants had no neurological, neuropsychological, or developmental diagnoses. The study was approved by the Massachusetts Institute of Technology and participants gave consent to participate.

##### Kids

Twenty-seven kids from the greater Boston area were used as part of a larger study. Kids were scanned after the completion of second grade (14 female, mean age = 8.4 years). All participants were born after at least 36 weeks gestation and had no neurological, neuropsychological, or developmental diagnoses. The study was approved by the Massachusetts Institute of Technology and Boston Children’s Hospital. Parents gave written consent and children gave verbal assent to participate.

##### Neonates

Neonatal data were acquired from the initial release of 40 neonatal subjects from the Developmental Human Connectome Project. Neonates were scanned at the Evelina Neonatal Imaging Center in London, and the study was approved by the UK Health Research Authority. The present study excludes data from neonates in which Freesurfer’s aparc+aseg anatomical segmentation was misaligned, resulting in 15 total participants. All neonates were born and at term age when imaged (of the 15: 5 female, mean gestational age at birth = 38.21, mean gestational age at scan = 39.05 weeks).

#### 2.1.2. Acquisition

For the adult and kid participants, diffusion-weighted data were acquired using echo planar imaging (64 slices, voxel size 2mm^3^, 128×128 base resolution, b-value 700s/mm^2^, diffusion weighting isotropically distributed along 60 directions) on a 3T Siemens scanner with a 32-channel head-coil. Images were processed using FSL’s FDT software. Additionally, a high-resolution (1 mm^3^) 3D magnetization-prepared rapid acquisition with gradient echo (MPRAGE) scan was also acquired on all participants. An online prospective motion algorithm reduced the effect of motion artifacts during the structural scan, and 10 selective reacquisition images were acquired and included to replace images that were affected by head motion. Structural MRI data were processed in FreeSurfer v.5.2.0 (http://surfer.nmr.mgh.harvard.edu/) using a semiautomated processing stream with default parameters, which includes motion and intensity correction, surface coregistration, spatial smoothing, subcortical segmentation, and cortical parcellation based on spherical template registration. The resulting cortical parcellations were reviewed for quality control.

For the neonate participants, the diffusion-weighted data used here were acquired using a spherically optimized set of directions on 4 shells with b-values of 0 (20 slices), 400 (64 slices), 1000 (88 slices), and 2600 (128 slices) (Tournier et al., 2015). These directions were split into four optimal subsets, one per phase encoding direction. The phase encoding directions were spread temporally, accounting for motion and duty cycle considerations. Additional motion correction reconstruction techniques were developed and interruption robustness was accomplished using a restart capability with user-selected set back from the break time point (Hutter et al., 2015, Hughes et al., 2017b). Slices were 3mm with 1.5mm overlap and 1.5×1.5mm resolution, with TE=90 and TR=3800ms. Additionally, a high-resolution (0.8 mm^3^) MPRAGE scan was also acquired on all participants. Images were acquired on a Philips 3T Achieva scanner using a specially designed neonatal 32 channel phased array head coil with dedicated slim immobilization pieces to reduce gross motion (Hughes et al., 2017b). Neonates were scanned while in natural sleep. Structural MRI data were processed in FreeSurfer v.6.0.0 (http://surfer.nmr.mgh.harvard.edu/fswiki/infantFS) using a dedicated infant processing pipeline (Zöllei et al., 2017; de Macedo Rodrigues et al., 2015; Makropoulos et al., 2018) which includes motion and intensity correction, surface coregistration, spatial smoothing, subcortical segmentation, and cortical parcellation based on spherical template registration. The resulting cortical parcellations were reviewed for quality control, and only neonates with correctly labeled cortical parcellations were used in the analysis.

#### 2.1.3. Defining seeds and targets for tractography

Using automated segmentation (Saygin et al., 2017), nine amygdala subnuclei (lateral, basal, accessory basal, central, medial, cortical, paralaminar, cortico-amygdaloid transition area, anterior amygdala area) were parcellated in each individual’s native space. In an attempt to replicate the gradient diagram from Freese and Amaral (2005, Figure 1) in humans, and because the lateral and basal subnuclei are associated with sensory and cognitive processes (Johansen et al., 2010; Stefanacci & Amaral, 2002) and thus likely contribute to emotional visual perception, the lateral and basal subnuclei were the two main seeds of interest in the present experiment (Figure 2A).

**Figure 2.**
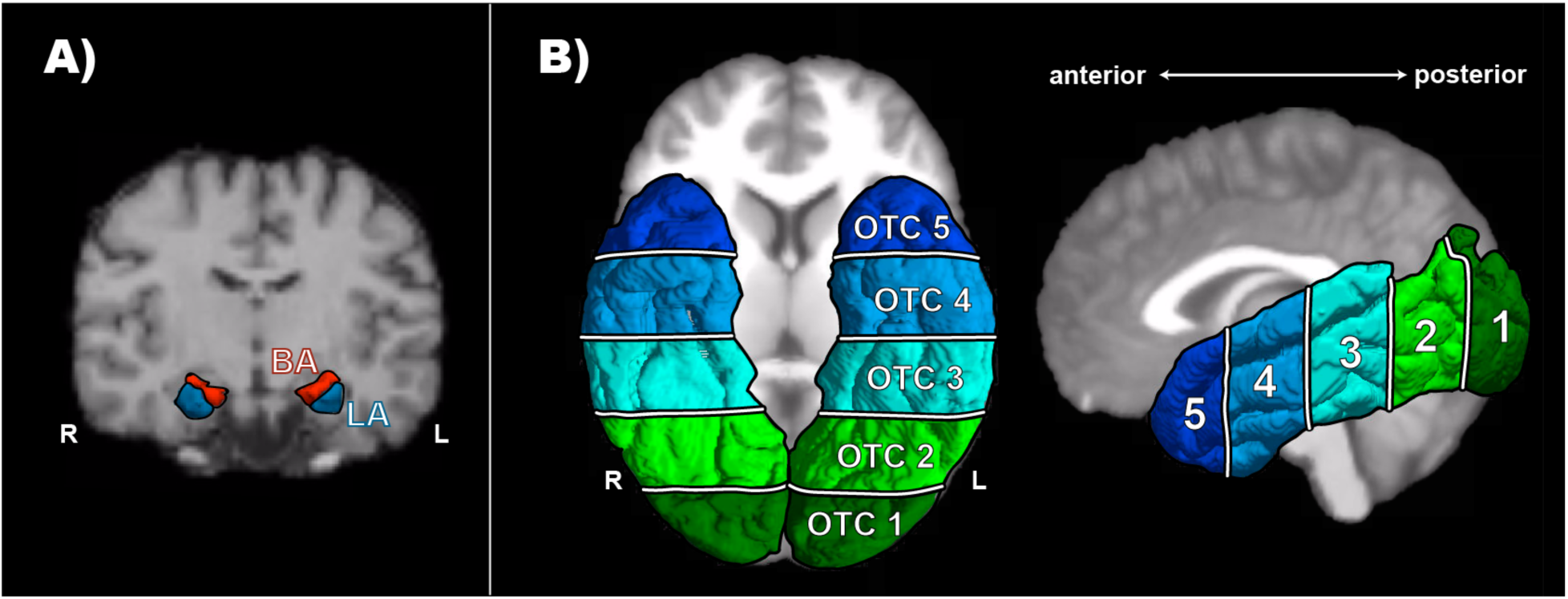
Seeds (A) and targets (B) used for tractography. A) Using the atlas developed by Saygin et al., 2017, an example parcellation of the lateral (LA, blue) and basal (BA, red) subnuclei in a representative subject, viewed coronally. B) Depiction of the 5 occipitotemporal cortex (OTC) labels in a representative subject viewed axially (left) and sagittally (right). Labels marked from most anterior (OTC 5, dark blue) to most posterior (OTC 1, dark green).

To explore the structural connectivity of the amygdala subnuclei to occipitotemporal cortex (OTC), an OTC label was made for each individual that combined all Freesurfer anatomical regions (using aparc+aseg) in the occipital and temporal cortices. In order to track differences in connectivity across the region, the label was split (separately for each individual and each hemisphere) into five equal sections from anterior to posterior (Figure 2B). These five OTC sections were the targets used for tractography.

Using the diffusion-weighted imaging data for each subject, principal diffusion directions were calculated per voxel. Probabilistic tractography was carried out using FSL-FDT (Behrens et al., 2003; Behrens et al., 2007; Tomassini et al., 2007) on the Ohio Supercomputer (Ohio Supercomputer Center, 1987) between the two amygdala subnuclei as seeds and five anatomical OTC sections as targets. Each seed voxel had 10,000 streamline samples to create a connectivity distribution to each of the target regions, while avoiding a mask consisting of the ventricles and cerebellum. The connection probability to each of the of the target regions was calculated for each amygdala voxel in each participant, producing a connectivity vector for each voxel. To account for differing connection probabilities in the deepest parts of the amygdala and to allow for relative comparison within and across participants (e.g., Croxson et al., 2005; Tomassini et al., 2007; Bach et al., 2011), each connectivity vector was normalized to [0,1] by dividing by the maximum connection probability of that amygdala voxel to all the other target regions (see Saygin et al., 2011).

### 2.2. Results and Discussion

Figure 3 depicts the connectivity that results from seeding the lateral subnucleus to the entire OTC, averaged across subjects in each sample, and projected onto a representative individual from each sample. Figure 4 depicts how the magnitude of this connectivity changes slice by slice in the entire OTC. A gradient-like decrease in connectivity was immediately apparent in the adults and kids, but neonates exhibited additional connectivity to the posterior cortex not found in the older samples.

**Figure 3.**
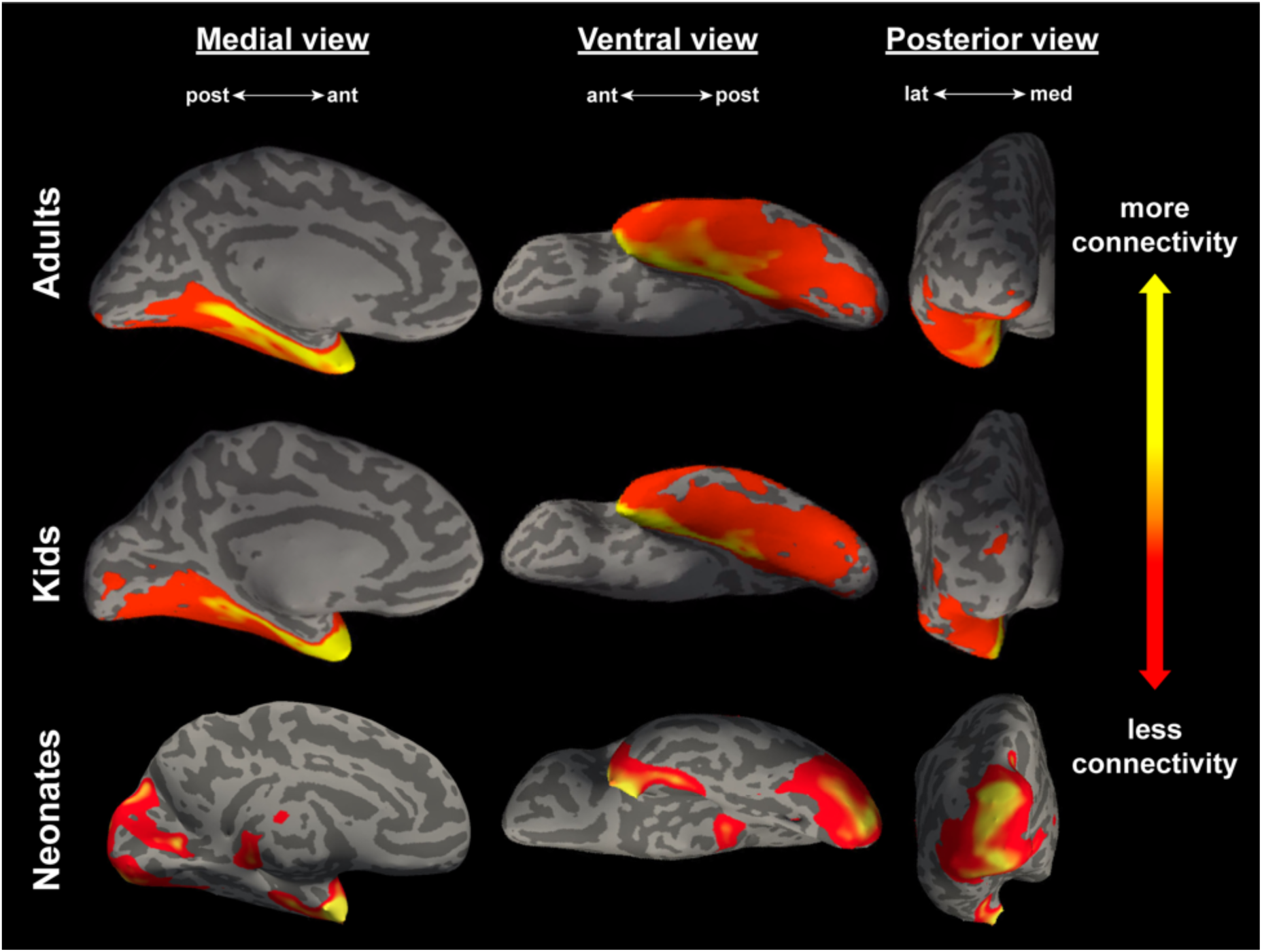
Connectivity between the lateral subnucleus and the entire occipitotemporal cortex, averaged across all adults (N = 20, top panel), kids (N = 27, middle panel), and neonates (N = 15, bottom panel). All views are of an inflated left hemisphere. Connectivity values are on a log scale thresholded for each sample such that yellow corresponds to a max percentile of 0.99, and red represents a min percentile of 0.85.

**Figure 4.**
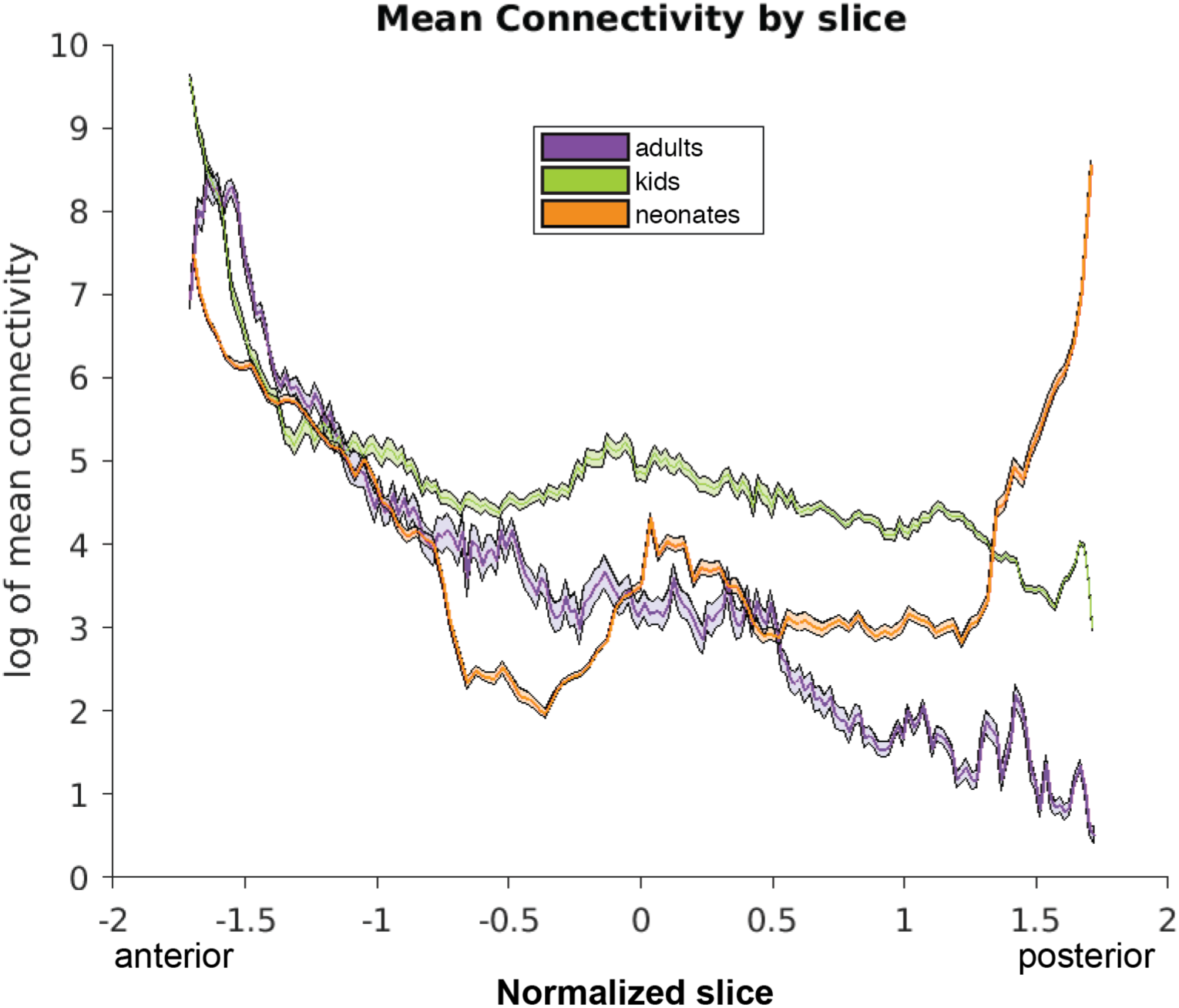
Connectivity between the lateral subnucleus and the entire occipitotemporal cortex, meaned across each slice in the OTC, moving coronally from anterior to posterior. Slices are normalized using z-scores of mean connectivity for each sample, thus accounting for differences in head size and allowing for comparison across samples. Lines represent the mean connectivity, shading represents standard error of the mean.

Further analyses were conducted by binning the OTC into the five sections used as targets for tractography. A 3 (Sample: adults vs. kids vs. neonates, between-subjects) × 2 (Amygdala subnucleus: lateral vs. basal, within-subjects) × 5 (OTC label: 5 [anterior] vs. 4 vs. 3 vs. 2 vs. 1 [posterior], within-subjects) ANOVA was conducted, to assess changes in mean connectivity from the amygdala subnuclei to the OTC across development. For analyses in which multiple comparisons were conducted, we used the Holm-Bonferroni method (Holm, 1979) to control the familywise Type I error rate (corrected p-values are denoted by *p*_HB_).

Results showed that indeed, across all seeds and samples and collapsed across hemispheres, there was a main effect of OTC label, *F*(4,590) = 559.25, *p* < 5 × 10^−199^ (Figure 5). More specifically, connectivity to each of the sections decreased on a gradient from anterior to posterior, with significantly more connectivity to OTC 5 than OTC 4 (*t*(123) = 17.01, *p*_HB_ < 2 × 10^−33^), to OTC 4 than OTC 3 (*t*(123) = 8.96, *p*_HB_ < 2 × 10^−14^), and to OTC 3 than OTC 2 (*t*(123) = 5.41, *p*_HB_ < 7 × 10^−7^). However, there was marginally more connectivity to OTC 1 than OTC 2 (*t*(123) = −1.94, *p*_HB_ = 0.054), although not significant.

**Figure 5.**
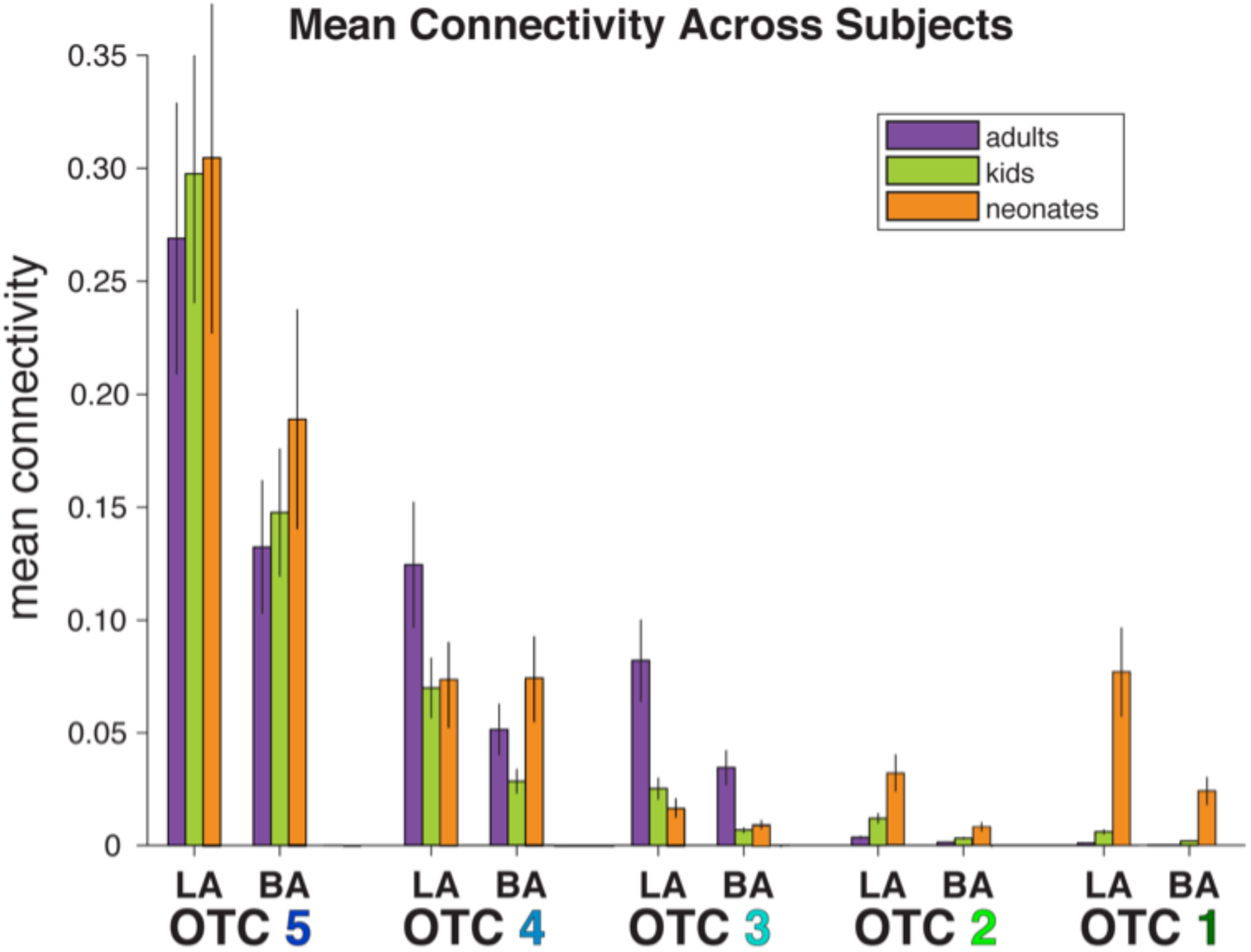
Mean connectivity, calculated separately in each individual and averaged across each sample, between the lateral (LA) and basal (BA) amygdala subnuclei and each of the five anatomical OTC labels. Error bars depict standard error of the mean.

This break in the gradient is reflected in a significant sample × OTC label interaction (*F*(8,590) = 10.67, *p* < 5 × 10^−14^), and as evidenced by Figure 5, can be attributed to the relatively higher connectivity to OTC 1 in neonates. In fact, when broken down by sample, adults and kids both show significant decreases across the entire OTC region (see Table 1). Neonates, however, display a U-shaped pattern: connectivity in neonates significantly decreases from anterior-to mid-OTC, then increases to posterior-OTC (Table 1). This suggests that all connections in kids and anterior connections in neonates are already adult-like since they follow a similar gradient pattern, but become more refined with age given significantly more connections to posterior regions in the neonates.

**Table 1.** t-test results and corresponding p-values comparing mean connectivity between each adjacent OTC label, collapsed across seed and broken down by sample.

| Sample | OTC comparison | *t* | *df* | *p* |
| --- | --- | --- | --- | --- |
| *Adults* | 5 – 4 | 8.467 | 39 | $9.118 \times 10^{-10}$ |
| | 4 – 3 | 3.870 | 39 | $8.058 \times 10^{-4}$ |
| | 3 – 2 | 7.568 | 39 | $1.080 \times 10^{-8}$ |
|  | 2 – 1 | 2.657 | 39 | 0.011 |
| *Kids* | 5 – 4 | 12.813 | 53 | $2.924 \times 10^{-17}$ |
| | 4 – 3 | 8.210 | 53 | $1.552 \times 10^{-10}$ |
|  | 3 – 2 | 3.332 | 53 | 0.003 |
|  | 2 – 1 | 2.706 | 53 | 0.009 |
| *Neonates* | 5 – 4 | 8.705 | 29 | $5.562 \times 10^{-9}$ |
| | 4 – 3 | 5.091 | 29 | $5.914 \times 10^{-5}$ |
|  | 3 – 2 | -1.570 | 29 | 0.127 |
|  | 2 – 1 | -3.229 | 29 | 0.006 |

Note: *p*-values are Holm-Bonferroni corrected.

Lastly, there was a significant main effect of mean connectivity between the two amygdala subnuclei seeds (*F*(1,590) = 183.83, *p* < 2 × 10^−36^), with the lateral subnucleus showing higher connectivity overall than the basal subnucleus, supporting previous research that each subnucleus has unique functions and connectivity patterns. Overall, the development of amygdalar connectivity to the OTC in humans resembles that of macaques, with certain exceptions that may correspond to differences in functional specialization of the OTC targets.

## 3. Experiment 2

Both macaques and humans exhibit a noteworthy pattern of connectivity between the lateral and basal subnuclei and OTC, and the connectivity changes are consistent across development between the two species. However, both of these findings probe connections to anatomical cortical targets. Further, Experiment 1 also revealed that neonates have significantly more connections to posterior cortex relative to kids and adults. Is there a functional explanation as to why the connectivity is organized in such a pattern? Or, more specifically, what functional regions in the OTC might be driving the developmental change in structural connectivity in humans revealed in Experiment 1?

Additionally, it is possible that an underlying scaffold exists within each parcel, matching the anatomical gradient, such that anterior sections of the parcels receive more connections than posterior sections. We therefore also divided up each parcel into anterior/posterior sections to address this possibility.

### 3.1. Method

#### 3.1.1. Participants/Acquisition

The same participants were used as in Experiment 1. The same diffusion scans (i.e., same acquisition parameters) as Experiment 1 were used in Experiment 2.

#### 3.1.2. Defining seeds and targets for tractography

As in Experiment 1, the lateral and basal amygdala subnuclei were the main seeds of interest. To explore what functional cortical regions in the OTC might be driving the gradient of connectivity observed by Freese and Amaral (2005) in Macaques and by Experiment 1 in humans, functional parcels were used as targets in Experiment 2. Functional parcels were chosen that were shown to encompass individually defined regions in the OTC associated with the visual processing of faces, scenes, and objects (Julian et al., 2012). These parcels included the Fusiform Face Area (FFA), Occipital Face Area (OFA), and Superior Temporal Sulcus (STS), regions known for processing faces over objects (e.g., Kanwisher, McDermott & Chun, 1997; Gauthier et al., 2000; Hoffman & Haxby, 2000); Lateral Occipital (LO), a subdivision of the Lateral Occipital Complex known for processing objects over scrambled objects (e.g., Grill-Spector et al., 1999); and Parahippocampal Place Area (PPA) and Retrosplenial Cortex (RSC), which process scenes over objects and are activated in tasks such as navigation (e.g., Epstein & Kanwisher, 1998; Epstein, 2008). Parcels were excluded if they overlapped more than 50% with another parcel; for this reason, we elected not to use the Posterior Fusiform Sulcus (PFS), the other subdivision of the Lateral Occipital Complex, due to its high overlap with FFA.

The functional selectivity of these parcels was validated in the adult subjects with four different localizer tasks. Once confirmed, the six parcels, originally in CVS average-35 MNI152 anatomical space, were then overlaid onto each individual’s anatomical MPRAGE scan using the inverse transform of Freesurfer’s CVS toolbox (Postelnicu et al., 2009; Zöllei et al., 2010; https://surfer.nmr.mgh.harvard.edu/fswiki/mri_cvs_register) to the CVS average-35 in MNI152 template in adults and kids, and Advanced Normalization Tools software (ANTs, Avants et al., 2011) in neonates. Then, the parcels were transformed to native diffusion space using nearest neighbor interpolation with Freesurfer’s bbregister function (Greve et al., 2009; https://surfer.nmr.mgh.harvard.edu/fswiki/bbregister) in adults and kids, and ANTs in neonates. It is worth noting that although neonates may not have functionally-specific cortical areas yet, registering the parcels into each individual’s native anatomy allows us to align the regions to a common space and approximate where the regions will eventually be located in neonates.

In addition to these six functional parcels, we sought to explore connectivity to primary visual cortex (V1); since the location of V1 is evolutionarily preserved, V1 was anatomically defined in each subject using the pericalcarine cortex.

Because some of these parcels overlap with each other, the seven targets were pruned by comparing each parcel against every other in each individual subject, locating any intersecting voxels, and assigning the voxels in question to whichever of the parcels was smaller; this eliminated ambiguity with tractography so that no voxel belonged to more than one target. Additionally, any white matter voxels contained within the parcels were also removed.

For continuity with Experiment 1, we wanted to determine the location of these seven parcels along the sagittal axis in order to sort them into OTC labels. This was done by loading each original parcel from the CVS average brain alongside the five OTC labels – which were anatomically defined in the CVS average brain as in Experiment 1 – and calculating the proportion of overlap between each parcel and each of the five labels. A given parcel was assigned to the OTC label with which it contained the highest proportion of overlapping voxels, thereby representing where most of the parcel’s mass was. These functional parcels are all located exclusively in the posterior half of the OTC (i.e., between OTC 1 and OTC 3); as such, we are not drawing functional conclusions about the anterior portions of the OTC. The functional parcels were sorted into the following OTC labels, collapsed across hemispheres: OTC 3 (STS, PPA), OTC 2 (RSC, FFA, LO), and OTC 1 (OFA,V1). See Figure 6A for parcel locations.

**Figure 6.**
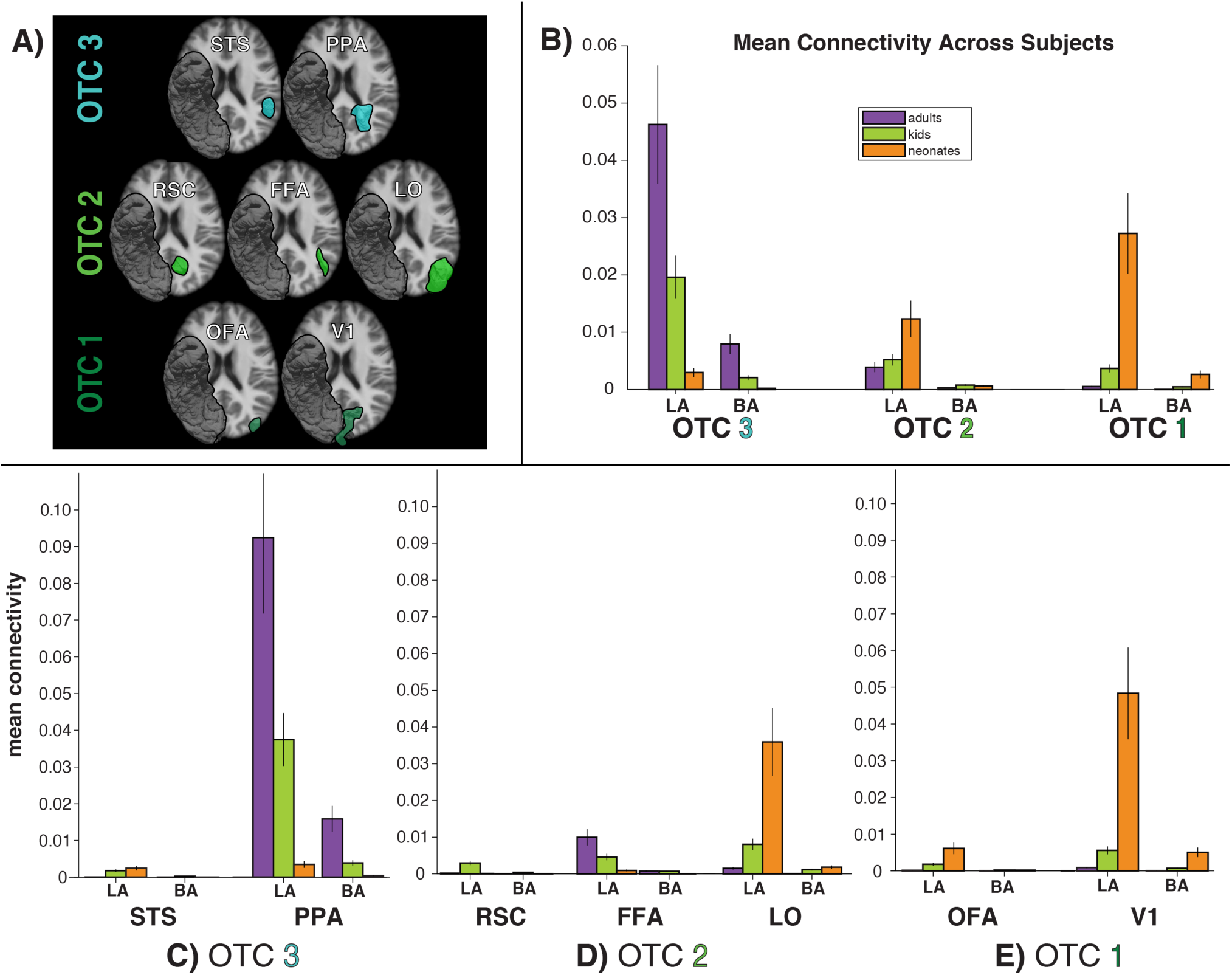
A) A 3D rendering of each of the seven functional parcels in the left hemisphere, viewed ventrally and overlaid on an axial slice. For reference, the right hemisphere contains the entire OTC. The seven parcels are sorted by their location, from anterior (OTC 3) to posterior (OTC 1). B) Mean connectivity, calculated separately in each individual and averaged across each sample, between the lateral (LA) and basal (BA) amygdala subnuclei and the three OTC labels that were the primary locations of functional parcels. Connectivity is collapsed across parcels: OTC 3 averages connectivity across STS and PPA. OTC 2 averages connectivity across RSC, FFA, and LO. OTC 1 averages connectivity across OFA and V1. C-E) Connectivity to the functional parcels individually, split depending on whether the parcel was located primarily in OTC 3 (C), OTC 2 (D), or OTC 1 (E). Error bars depict standard error of the mean.

As before, using the diffusion-weighted imaging data for each subject, probabilistic tractography was run with FSL, this time with the amygdala subnuclei as seeds and functional parcels as targets (using the same procedures as in Exp. 1)

Additionally, in a separate analysis exporing an underlying connectivity scaffold within the parcels, the seven functional parcels were divided in half in native diffusion space, creating separate anterior and posterior sections for each parcel. Probabilistic tractography was again run with FSL, using the amygdala subnuclei as seeds and these anterior and posterior subdivisions as targets.

### 3.2. Results and Discussion

First, a 3 (Sample: adults vs. kids vs. neonates, between-subjects) × 2 (Amygdala subnucleus: lateral vs. basal, within-subjects) × 3 (OTC label: 3 [anterior] vs. 2 vs. 1 [posterior], within-subjects) ANOVA was conducted to assess developmental changes in mean connectivity from the amygdala subnuclei to functional parcels located in the OTC. If these functional parcels were at least somewhat contributing to the gradient found in Experiment 1, then results should be similar to the anatomical findings; if the connectivity to anatomical targets is independent of the functional specialization of the cortex, then a different pattern might emerge.

Results showed that indeed, across all seeds and samples and collapsed across hemispheres, there was a main effect of OTC label, *F*(2,354) = 22.98, *p* < 5 × 10^−10^ (Figure 6B). OTC 3 had significantly higher mean connectivity that OTC 2 (*t*(123) = 5.14, *p*_HB_ < 4 × 10^−6^) and OTC 1 (*t*(123) = 3.83, *p*_HB_ < 5 × 10^−4^). However, connectivity differences were not significantly different between OTC 2 and OTC 1 (*t*(123) = −1.11, *p*_HB_ = 0.27) when collapsed across samples. As evidenced by the graphs of mean connectivity in Figure 5 and Figure 6B, these results mirror the anatomical findings from Experiment 1. This confirms that the functional parcels used here contributed at least somewhat to the observed anatomical gradient. To discern which functional parcels were driving the gradient, additional analyses were done comparing the connectivity to the parcels located within each OTC label separately.

In OTC 3 (Figure 6C), collapsing across seeds and samples, connectivity was significantly higher to PPA than it was to STS (*t*(123) = −6.71, *p* < 7 × 10^−10^). Within PPA, connectivity was significantly higher in adults than kids (*t*(92) = 3.65, *p*_HB_ < 5 × 10^−4^) and in kids than neonates (*t*(82) = 4.14, *p*_HB_ < 2 × 10^−4^). This suggests that the connectivity to the middle of the OTC is largely driven by connections to PPA (a region known for processing scenes) relative to STS, and that this connectivity increases across development.

In OTC 2 (Figure 6D), LO had significantly more connectivity from the lateral and basal subnuclei, collapsed across samples, compared to both FFA (*t*(123) = −2.15, *p*_HB_ = 0.03) and RSC (*t*(129) = −3.71, *p*_HB_ < 1 × 10^−3^). Within LO, connectivity was significantly lower in adults than kids (*t*(92) = −2.62, *p*_HB_ = 0.01) and in kids than neonates (*t*(82) = −3.09, *p*_HB_ = 0.005). This suggests that connectivity to mid-to-posterior OTC is largely driven by connections to LO (a region known for processing objects) more than it is by connections to FFA or RSC, and that these connections to LO decrease across development.

In OTC 1 (Figure 6E), V1 had significantly more connectivity from the lateral and basal subnuclei than did OFA, collapsed across samples and hemispheres (*t*(123) = −3.66, *p* < 4 × 10^−4^). Within V1, connectivity was marginally lower in adults than kids (*t*(92) = −1.88, *p*_HB_ = 0.063) and significantly lower in kids than neonates (*t*(82) = −4.03, *p*_HB_ < 4 × 10^−4^). This suggests that connectivity to posterior OTC is largely driven by connections to primary visual cortex, more so than by connections to OFA, and that this connectivity decreases across development.

Further, we explored how connectivity was distributed across the functional parcels by looking at anterior and posterior divisions separately, using a 3 (Sample: adults vs. kids vs. neonates, between-subjects) × 2 (Amygdala subnucleus: lateral vs. basal, within-subjects) × 2 (subdivision: anterior vs. posterior, within-subjects) ANOVA. Connectivity differences between the parcel subdivisions are depicted in Supplementary Figure S1. Noteably, there was a significant sample × subdivision interaction (*F*(2,1652) = 23.28, *p* < 2 × 10^−10^), as well as a significant parcel × subdivision interaction (*F*(6,1652) = 35.28, *p* < 3 × 10^−40^). Collapsed across seeds and parcels, adults and kids had significantly higher connectivity to the anterior subdivisions (adults: *t*(279) = 4.73, *p*_HB_ < 8 × 10^−6^, kids: *t*(377) = 5.23, *p*_HB_ < 9 × 10^−7^), whereas neonates had marginally higher connectivity to posterior subdivisions (*t*(209) = −1.64, *p*_HB_ = 0.102). For simplicity, we report results for the three parcels determined above to have the highest contribution to the observed connectivity: PPA, LO, and V1.

Within PPA, collapsing across seeds and samples, connectivity was significantly higher to the anterior half than to the posterior half (*t*(123) = 6.35, *p*_HB_ < 3 × 10^−8^). There was a significant sample × subdivision interaction (*F*(2,236) = 27.18, *p* < 3 × 10^−11^); adults and kids had significantly higher connectivity to the anterior section (adults: *t*(39) = 5.73, *p*_HB_ < 4 × 10^−6^, kids: *t*(53) = 5.16, *p*_HB_ < 8 × 10^−6^), whereas neonates had marginally higher connectivity to the posterior section (*t*(29) = −2.01, *p*_HB_ = 0.054).

Within LO, collapsing across seeds and samples, connectivity was not significantly different between the anterior half and the posterior half (*t*(123) = 1.95, *p*_HB_ = 0.266). There was also no sample × subdivision interaction (*F*(2,236) = 0.88, *p* = 0.416).

Within V1, collapsing across seeds and samples, connectivity was likewise not significantly different between the anterior half and the posterior half (*t*(123) = −2.17, *p*_HB_ = 0.127). However, there was a significant sample × subdivision interaction (*F*(2,236) = 6.65, *p* = 0.002); neonates had marginally higher connectivity to the posterior section (*t*(29) = −2.27, *p*_HB_ = 0.092), whereas adults and kids had no significant differences.

Overall, analysis of the parcel subdivisions suggests that the functional specialization of the posterior OTC is equally distributed across the entirety of the most connected parcels, whereas connectivity to functional parcels in the mid-OTC has more of an anterior bias. Moreover, the connectivity distributions of these parcels seem to undergo a developmental shift, from more distributed/posterior to more anterior, after early development.

Lastly, as in Experiment 1, there was a significant main effect of mean connectivity between the two amygdala subnuclei seeds, in both the whole parcel analysis (*F*(1,354) = 99.72, *p* < 8 × 10^−21^) and in the subdivided parcel anaylsis (*F*(1,1652) = 132.97, *p* < 2 × 10^−29^). The lateral subnucleus showed higher connectivity overall than the basal subnucleus, supporting again that the lateral and basal subnuclei have unique functions and connectivity patterns.

## 4. General Discussion

Investigating the structural connectivity between the amygdala and occipitotemporal cortex will help us better understand the amygdala’s role in perceiving and processing emotional visual stimuli, which has ecological relevance and certainly changes across development. Many functionally specialized visual regions exist within occipitotemporal cortex, but it was previously unknown how connectivity to these regions develops from birth in humans. Previous work in macaques had revealed connections between the lateral and basal amygdala subnuclei and the occipitotemporal cortex, noting a rostrocaudal topographic organization of the connections (e.g., Freese & Amaral, 2005) and refinement across development (e.g., Webster et al., 1991a). In this paper, we explored this topographic organization in humans using noninvasive methods, and further investigated specific functional cortical areas located within the occipitotemporal region that may be contributing to the observed pattern of structural connectivity.

In our study, connectivity between the lateral and basal subnuclei and occipitotemporal cortex in human adults significantly decreased on a gradient, replicating the finding in macaques. Likewise, kids displayed a significantly decreasing gradient, upholding previous conclusions that connections are already adult-like early in development. However, one notable result in our findings was an almost U-shaped trend in neonates: although neonates had significantly decreasing connectivity moving from anterior to middle occipitotemporal cortex, connectivity significantly increased again to posterior cortex. These results support previous work in animals (e.g., Webster et al., 1991a; 1991b) and humans (Saygin et al., 2015; Gabard-Durnam et al., 2018) that infants and kids show similar patterns of connectivity as adults, as well as similar functional organization of visual cortex (Deen et al., 2017), with additional refinement and pruning that occurs through typical development after relevant cognitive milestones and functional specialization occur (Gogtay et al., 2004; O’Leary, 1992; Khundrakpam et al., 2013).

Splitting the cortex into functionally defined parcels allowed us to further hone in on the developmental changes in this pattern. We found that particular regions, specifically PPA, LO, and V1, contribute heavily to the refinement of connectivity to mid-to posterior occipitotemporal cortex. Our results match what is known for adults, for example that RSC and the occipital lobe do not project to the amygdala (Stefanacci & Amaral, 2000; Stefanacci & Amaral, 2002), that STS has very weak projections (Stefanacci & Amaral, 2000), and that the parahippocampal cortex has high connections (Saygin et al., 2015) with the adult amygdala.

Further analyses exploring anterior and posterior subdivisions of the functional regions revealed that connections to PPA are especially driven by the anterior portion of the parcel, whereas connections to LO and V1 are distributed across the entire parcel. This finding is again supported by tracer studies, for example from Freese and Amaral (2005), which noted that most amygdaloid fibers that extend to area TE in macaques enter gray matter at more rostral levels of area TE, whereas amygdaloid fibers to area V1 are evenly distributed throughout the rostrocaudal and dorsoventral extent of the region. Alternatively, the subdivision analyses presented here may elucidate what part of the parcel likely represents the location of the true functional region in each subject: perhaps the cortex containing a subject’s actual PPA is more anterior, whereas a subject’s LO is better captured by the entire parcel.

As a whole, each of these parcels exhibited noteworthy changes in connectivity across development. Adults and kids exhibit significantly higher connectivity to PPA (a region known for processing scenes) and neonates exhibit drastically higher connectivity to both LO (a region known for processing objects) and primary visual cortex. Given that these neonates were scanned soon after birth, visual input was still novel, visual acuity was still poor, and experience with real-world objects was lacking. Thus, we speculate that this is why V1 shows the biggest developmental trend in our data; perhaps V1 is strongly influenced by the amygdala in processing emotional visual stimuli at birth, and thus has more connections to the amygdala than is necessary later in development.

Other work exploring anatomical and functional cortical maturation may additionally explain the developmental trends in our results. For instance, it has been shown in a longitudinal study of 4- to 21-year-olds that occipitotemporal cortex matures differentially across development, with occipital and temporal poles maturing earliest and the rest of the temporal cortex maturing later, after memory association areas develop (Gogtay et al., 2004). This may inform the relatively high connections we found in neonates to the most anterior and posterior sections of the occipitotemporal cortex, given that these cortical areas are the first to mature and memory systems are immature in infants. It is quite possible that regions like PPA and LO are continuing to mature through infancy (e.g., Deen et al.) and that this functional maturation is contributing to the maturational changes in connectivity that we find here. Future work is needed to explore the age at which this transition happens, and how soon after birth maturation and pruning occurs.

The developmental origins of amygdalar-occipitotemporal connectivity may offer additional insight into how human cortex develops its specificity and functional organization. It has been shown that connectivity is largely established at birth: structural connectivity between the amygdala and temporal regions already exists in utero, by 13 weeks post-conception (Vasung et al., 2010). However, as evidenced by the present experiments, although the same connections are largely in place at birth, activity-dependent interactions between cortical regions may fine-tune their functionality. Experience with the environment postnatally may lead to notable age-related changes. The functional maturation of the occipitotemporal regions that were studied in the present experiments may contribute to the continued refinement of the amygdalar subregions, the amygdalar connections to these regions, and to the functional specialization of occipitotemporal regions themselves (e.g. in line with Interactive Specialization theories of development; Johnson 2010).

Additionally, the present experiments contribute to our understanding of the functional specification of the amygdala subnuclei. As noted previously, the human amygdala can be subdivided into at least nine unique subnuclei with distinct connectivity and functional patterns; here, we focused on the lateral and basal subnuclei, given their involvement with visual processing and the high connections between the basolateral subunit and occipitotemporal regions found in previous studies – but now at an unprecedented finer level than what was previously studied in humans. In both Experiments 1 and 2, we observed significant main effects of subnucleus: lateral had significantly higher connectivity to the occipitotemporal anatomical and functional targets than basal did. These results are concordant with the results of Stefanacci and Amaral (2000; 2002), who used lesion experiments in macaques to investigate the roles of cortical inputs to the amygdala subnuclei and hypothesized the functional significance of these inputs. They proposed that the primary function of the lateral subnucleus is to receive visual sensory input and transmit this information to the basal subnucleus. The basal subnucleus then integrates this sensory input with contextual information, via connections with the orbitofrontal and medial temporal cortices, and evaluates whether a “fear response” is necessary (Stefanacci & Amaral, 2002). Our results in humans corroborate this hypothesis and support the functional distinction between lateral and basal, given that the lateral subnucleus was more connected to visual sensory areas than the basal subnucleus across all stages of development.

One noteable limitation of the present study is that we use functional parcels originally defined in adults, and recognize that overlaying them onto neonates may have the potential to overestimate or mischaracterize certain cortex. However, we use ANTs to register the functional parcels to each neonate’s native space, which has been shown to be highly effective and reliable (Tustison et al., 2014). These limitations can only be overcome by scanning participants from all samples within the same study, and by functionally defining regions of interest individually, which may not be reliable in a sample of newborns (see Batalle et al., 2018 for a review of neuroimaging methods in adults vs. neonates).

Overall, the present experiments make apparent a decreasing pattern of structural connectivity between the amygdala subnuclei and occipitotemporal cortex, evidence for which has been repeatedly shown in macaques but was otherwise lacking in humans. Further, we use multiple cross-sectional samples to gauge the progression of this connectivity across development, and offer specific functional cortical areas in the ventral visual stream that might be driving the observed pattern of connectivity changes. This work has important clinical applications: Given the role of the amygdala in many psychiatric disorders – many of which have early onsets, such as autism and anxiety (e.g., Baron-Cohen et al., 2000; Lonigan & Phillips, 2001; Pine, 2007; Qin et al., 2014; Warnell et al., 2018) – it is crucial to fully understand how the amygdala connects to the rest of the brain across early development. Even more so, given that the amygdala subnuclei are distinct in their functions and connections, studying the amygdala at this finer level can elucidate even more information than would otherwise be gotten from treating the amygdala as a whole contiguous structure or as larger subunits. The developmental progression of structural connectivity between the amygdala subnuclei and occipitotemporal cortex in typically-developing humans can help us better understand developmental disorders or deficits implicated when these connections are abnormal or lacking. Further research can seek to explore this connectivity in patient populations, classify differences between patients and controls, and offer new diagnostic or treatment interventions.

### 4.1. Conclusions

The present experiments use DWI data to explore the structural connectivity between the lateral and basal subnuclei of the amygdala and the occipitotemporal cortex, which has been studied extensively in macaques but is otherwise lacking in humans. We first replicate in adult humans the connectivity gradient observed in adult macaques, and additionally show the developmental progression of this connectivity using cross-sectional samples of 8-year old kids as well as neonates. Additionally, we investigate a functional explanation for the observed anatomical gradient by targeting functionally defined parcels in the ventral visual stream, followed by an even finer-grained approach by splitting the parcels into anterior and posterior subsections. Our results suggest that, although connectivity between the amygdala subnuclei and occipitotemporal cortex in humans is largely adult-like at birth, the functional maturation of connections to anterior PPA, LO, and V1 seem to be driving the developmental changes that do occur.

## Acknowledgements

Support was provided by the Alfred P. Sloan Foundation and NICHD/NIH grant F32HD079169 (to Z.M.S). Analyses were completed using the Ohio Supercomputer. We would like to thank Nancy Kanwisher and other members of the Kanwisher lab for help with data acquisition, David Osher for comments and suggestions, and members of the Saygin Developmental Cognitive Neuroscience Lab for feedback and comments.

## Supplementary

**Figure S1.**
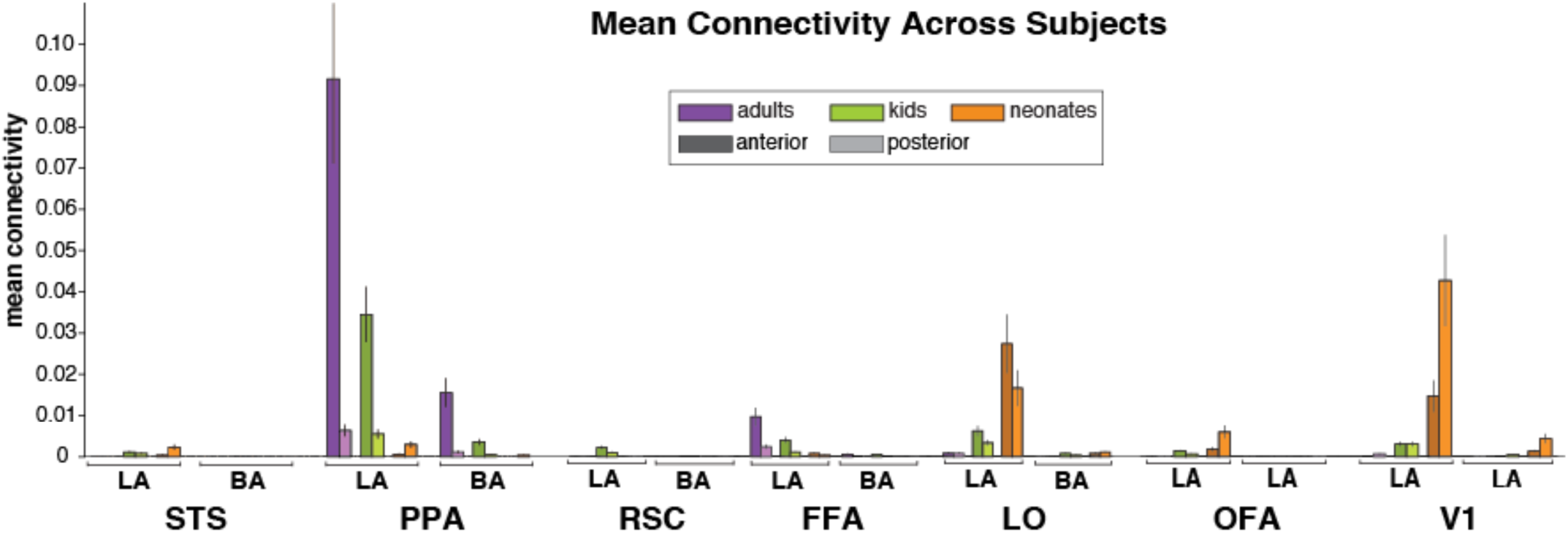
Mean connectivity from the lateral (LA) and basal (BA) subnuclei to the anterior (dark bars) and posterior (light bars) subdivisions of each functional parcel, within each sample. Error bars depict standard error of the mean.

